# Role of desmoplakin in supporting neuronal activity, neurogenic processes, and emotional-related behaviors in the dentate gyrus

**DOI:** 10.1101/2023.11.17.567646

**Authors:** Keisuke Otsubo, Naoko Sakashita, Yuki Nishimoto, Yo Sato, Takehisa Tsutsui, Katsunori Kobayashi, Kanzo Suzuki, Eri Segi-Nishida

## Abstract

Desmoplakin (Dsp) is a component of desmosomal cell-cell junctions that interacts with the cadherin complex and cytoskeletal intermediate filaments. In addition to its function as an adhesion component, Dsp is involved in various biological processes, such as gene expression, differentiation, and migration. Dsp is specifically expressed in the hippocampal dentate gyrus (DG) in the central nervous system. However, it is unclear how Dsp impacts hippocampal function and its related behaviors. Using an adeno-associated virus knockdown system in mice, we provide evidence that Dsp in the DG maintains hippocampal functions, including neuronal activity and adult neurogenesis, and contributes to anxiolytic-like effects. Dsp protein is mostly localized in mature granule cells in the adult DG. Dsp knockdown in the DG resulted in a lowered expression of an activity-dependent transcription factor FosB, and an increased expression of mature neuronal markers, such as calbindin. In addition, the suppression of Dsp decreases serotonin responsiveness at the DG output mossy fiber synapses and alters adult neurogenic processes in the subgranular zone of the DG. Moreover, DG-specific Dsp knockdown mice showed an increase in anxiety-like behaviors. Taken together, this research uncovers an unexplored function for Dsp in the central nervous system and suggests that Dsp in the DG may function as a regulator to maintain proper neuronal activation and adult neurogenesis, and contribute to the adaptation of emotion-related behavior.

## Introduction

Desmosomes are adhesive intercellular junctions that provide strong adhesion between cells and are abundant in tissues exposed to mechanical stress, such as the heart and skin. Mutations in these factors cause skin and heart vulnerability (Kowalczyk and Green, 2013). Desmosomal components comprise transmembrane cadherins, such as desmoglein and desmocollin, and intracellular proteins, such as plakoglobin, plakophilin, and desmoplakin (Dsp) (Garrod and Chidgrey, 2008; Broussard et al., 2015). Dsp is a family of plakin proteins that interacts with the cadherin/plakoglobin/plakophilin complex through the N-terminal domain and with an intermediate filament cytoskeleton through the C-terminal domain (Delva et al., 2009). In addition to mediating cell-to-cell adhesion, desmosomes function as cell signaling regulators involved in cell proliferation and differentiation (Johnson et al. 2014). In particular, Dsp is involved in processes such as gene expression, differentiation, and microtubule dynamics, indicating its diverse biological functions in addition to its function as an adhesion component (Bendrick et al., 2019; Dubash et al., 2016; Lombardi et al., 2016; Patel et al., 2014).

In the central nervous system, Dsp is specifically expressed in the hippocampal dentate gyrus (DG) in the brain (Yamasaki et al., 2008). The DG gates the flow of neocortical information into the hippocampal CA3 region and is one of the few areas of the mammalian brain where adult neurogenesis occurs (Drew et al., 2013; Zhao et al., 2008). It has been reported that Dsp is characterized as a molecular maker for neuronal maturation in the DG of hippocampus (Kobayashi et al., 2010). The expression level of *Dsp* in the DG increases with postnatal development, similarly to the patterns of other DG maturation markers, calbindin and tryptophan-2,3-dioxygenase 2 (Tdo2). Dsp transcription is directly regulated by Bcl11b, a transcription factor that plays an important role in postnatal development of the DG (Simon et al., 2012). Forebrain neuron-specific knockout of Dsp in Emx1-Cre mice impairs progenitor proliferation and neuronal differentiation at 14 days postnatal age, indicating that Dsp contributes to the development of dentate granule neurons (Simon et al., 2012). However, whether Dsp regulates the function of dentate mature granule neurons and affects the process of adult neurogenesis in the DG remain unclear.

The hippocampus is a limbic structure implicated in the regulation of stress and pathophysiology/treatment of anxiety and depression (Segi-Nishida and Suzuki, 2022). We have previously demonstrated that long-term antidepressant treatment suppresses the expression of *Dsp* in the DG (Kobayashi et al., 2010; Imoto et al., 2017), indicating that Dsp plays a key role in emotion-related behaviors in the DG. Nevertheless, whether Dsp in the DG during adulthood is involved in emotion-related behaviors such as depression and anxiety is unclear.

In this study, we investigated the effect of Dsp knockdown in the DG on neuronal activation, adult hippocampal neurogenesis, and emotion-related behaviors using an adeno-associated virus (AAV) knockdown system in mice. We found that Dsp in the adult hippocampal DG maintains the function of dentate granule neurons, regulates neuronal differentiation during adult neurogenesis, and behaviorally contributes to anxiolytic-like effects.

## Materials and Methods

### Experimental animals

Seven-to-eight-week-old male C57BL/6N mice (23–25 g) were purchased from Japan SLC (RRID5295404; Hamamatsu, Japan). The mice were housed in groups of six per cage (32 cm length × 11 cm width × 13.5 cm height). All the mice were housed under standard conditions (temperature: 24 ± 2 °C, humidity: 55% ± 5%) with a 12 h light/dark cycle and *ad libitum* access to water and food (MR standard, Nosan Corporation, Yokohama, Japan). The weight gain of each mouse was recorded during the experiments to monitor their physical condition. All mice were habituated for longer than one week before the experimental procedures were performed. A total of 132 mice were used in this study (control animals, n=14; ECS-treated animals, n=6; fluoxetine-treated animals, n=7; AAV control miRNA infected animals, n=52; AAV Dsp miRNA infected animals, n=53). Animal use and procedures were conducted in according the Tokyo University of Science regulations on animal care and use in research and were approved by the Animal Care and Use Committee of Tokyo University of Science (approval numbers K20009, K21007, and K22007).

### AAV preparation and hippocampal administration

AAVs (serotype rh10) expressing artificial microRNAs (miRNAs) targeting Dsp or control miRNAs with Emerald Green Fluorescent Protein (EmGFP) under the EF1α promoter were generated as follows: We designed the sequences for artificial microRNAs for Dsp according to the BLOCK-iT RNAi Designer website (Thermo Fisher Scientific). First, the DNA fragment encoding miRNA for *Dsp* (5’-TGC TGT TGA GAT GGA CAT CAA TGC ACG TTT TGG CCA CTG ACT GAC GTG CAT TGG TCC ATC TCA A-3’ and 5’-CCT GTT GAG ATG GAC CAA TGC ACG TCA GTC AGT GGC CAA AAC GTG CAT TGA TGT CCA TCT CAA C-3’) was subcloned into pcDNA6.2-GW-EmGFP-miR using BLOCK-iT Pol II miR RNAi Expression Vector Kit (Invitrogen). The EmGFP and Dsp miRNA fragments were amplified by PCR from this vector and inserted between the KpnI and EcoRV sites of the pAAV-EF1α-DIO EYFP vector (#27056; Addgene). The resulting plasmid was named pAAV-EF1α-EmGFP-miR-Dsp. The DNA fragment encoding control microRNA (5′-TGC TGA AAT GTA CTG CGC GTG GAG ACG TTT TGG CCA CTG ACT GAC GTC TCC ACG CAG TAC ATT T-3′ and 5′-CCT GAA ATG TAC TGC GTG GAG ACG TCA GTC AGT GGC CAA AAC GTC TCC ACG CGC AGT ACA TTT C-3′), which can form a hairpin structure processed into mature microRNA but is predicted not to target any known vertebrate gene, was used to generate an AAV vector expressing control microRNA. TheBLOCK-iT Pol II miR RNAi Expression Vector Kit was used instead of those encoding Dsp miRNA. AAV particles were produced using the previously described procedure with minor modifications (Shinohara et al., 2018). Briefly, the AAVrh10 expression plasmid (Addgene #112866), pHelper vector (#240071, Agilent Technologies, Santa Clara, CA), and the constructed AAV-vector plasmid were transfected into AAV-293 cells (#240073, Agilent) using Lipofectamine 2000 (Thermo Fisher Scientific). Transfected AAV-293 cells were collected 72 h after transfection and resuspended in artificial cerebrospinal fluid. The cell suspensions were freeze-thawed three times. Cell debris was eliminated by centrifugation at 10,000 × g for 10 min and the supernatant containing the virus was collected. The viral suspension was incubated with benzonase (Merck Millipore, Burlington, MA, USA) to degrade residual DNA. The prepared AAV solution was aliquoted and stored at −80 °C. The AAV titer was quantified using PCR (5′-TGA GTC ACC CAC ACA AAG GA-3′ and 5′-CCA AGC TGG CCT AAC TTC AG-3′) after proteinase K treatment (Merck Millipore). Under anesthesia with a mixture of medetomidine (0.3 mg/kg, ZENOAQ, Japan), midazolam (4 mg/kg, Sandoz, Japan), and butorphanol (5 mg/kg, Meiji, Japan), the mouse brain was fixed with stereotaxic instruments (Narishige, Tokyo, Japan). An AAV solution of 500 nL per injection site (5.0 × 10^8^ copies) was bilaterally injected into the DG using a PV-820 Pneumatic PicoPump (World Precision Instruments, Hessen, Germany) through a glass micropipette made with a PN-30 micropipette puller (Narishige). The stereotaxic coordinates were targeted to the ventral DG: 3.2 mm posterior to the bregma, 2.7 mm lateral to the midline, and 3.0 mm ventral to the skull surface at the bregma, according to the mouse brain atlas (Paxinos and Franklin, 2012). Data were excluded from the analysis if fluorescence was not observed in the DG.

### Electroconvulsive stimulation and drug treatment

Bilateral ECS (current, 25–35 mA; shock duration, 1 sec; frequency, 100 pulse/sec; pulse width, 0.5 msec) was administered with a pulse generator (ECT Unit; Ugo Basile, VA, Italy) to mice as described previously (Imoto et al., 2017). ECS was administered 4 times a week. Fluoxetine hydrochloride (LKT Labs, St. Paul, MN, USA) was dissolved in drinking water and orally applied at a dose of 22 mg/kg/day for 4 weeks as described previously (Kobayashi et al., 2010).

### RNA extraction and real-time PCR

Seven to eight weeks after AAV infection, the mice were decapitated under deep anesthesia, coronal brain slices (1 mm) were cut using a tissue slicer, and the DG of the hippocampus was dissected. Total RNA was extracted using the Reliaprep RNA Cell Miniprep System (Promega, Madison, WI, USA) and reverse transcribed using ReverTra Ace (Toyobo, Osaka, Japan). Subsequent, real-time PCR was performed using the StepOne system (Applied Biosystems, Foster City, CA) and Thunderbird SYBR qPCR mix (Toyobo). The expression levels of each gene were quantified using standardized external dilutions. The relative expression levels of the target genes were normalized to those of 18S rRNA or *Gapdh*. The specificity of each primer set was confirmed by melt-curve analysis, and the product size was examined using gel electrophoresis. Table 1 summarizes the primer sequences for each gene.

**Table 1.**
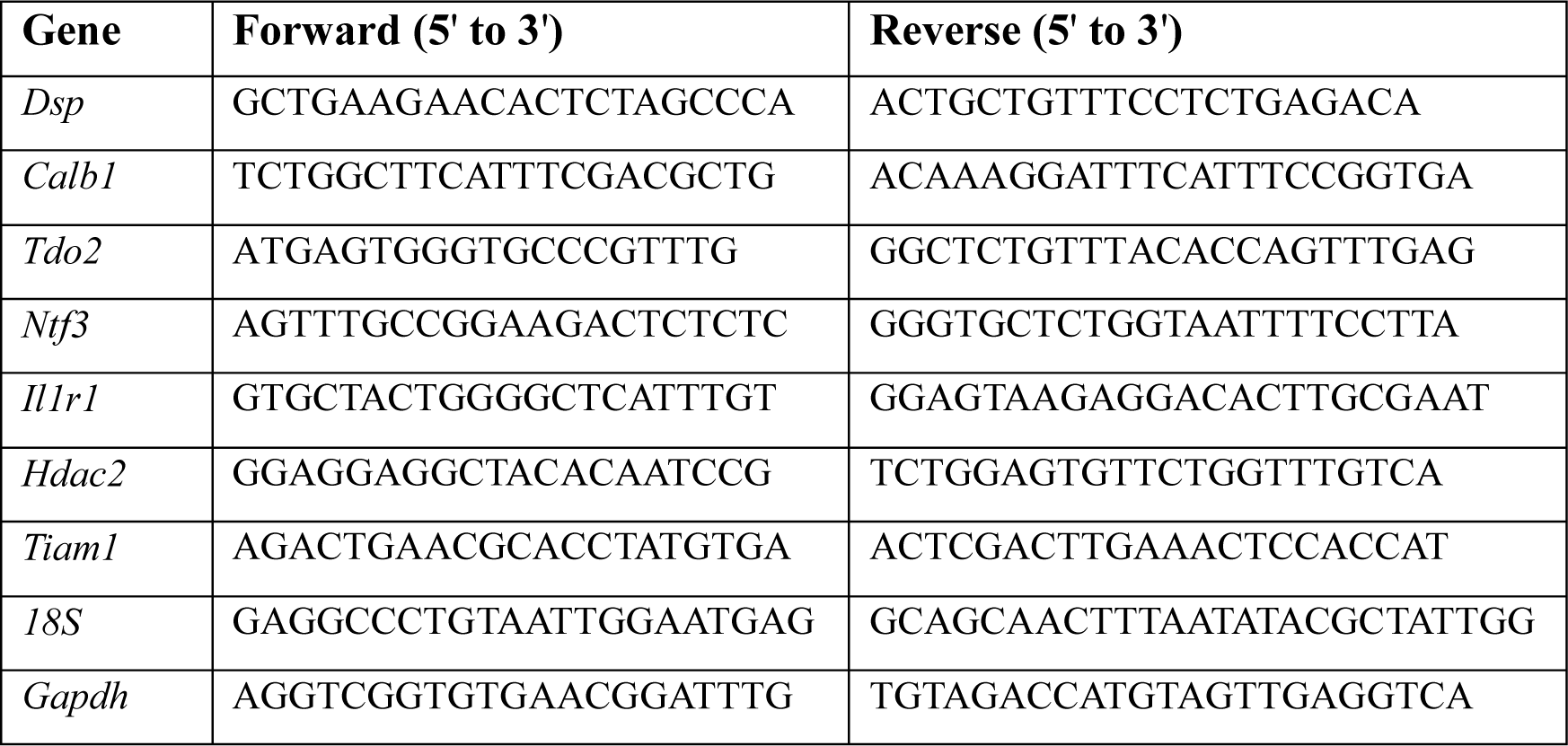
List of primers used for qPCR analysis.

### Electrophysiological analysis

Mice were decapitated under deep anesthesia 7–8 weeks after AAV infection, and both hippocampi were isolated. Transverse hippocampal slices (380 μm) were cut in ice-cold saline using a tissue slicer (7000smz, Campden Instruments Ltd., Leics, UK). The slices were then incubated for 30 min at 30 °C and maintained in a humidified interface-holding chamber at room temperature before use. Intense GFP fluorescence in the DG and the mossy fiber (MF) tract was confirmed, and electrophysiological recordings were performed as described (Kobayashi et al., 2010, 2020). Electrophysiological recordings were made in a submersion-type chamber maintained at 27.0–27.5 °C and superfused at 2 mL/min with recording saline composed of (in mM): NaCl, 125; KCl, 2.5; NaH2PO4, 1.0; NaHCO3, 26.2; glucose, 11; CaCl2, 2.5; MgCl2, 1.3 (equilibrated with 95% O2 / 5% CO2). For recording field excitatory postsynaptic potentials (EPSPs) arising from the medial perforant path (MPP)-granule cell synapse, a glass recording pipette filled with 2 M NaCl and bipolar tungsten stimulating electrodes were placed in the middle third of the molecular layer in the dentate gyrus. The input output relationship at the MPP input was examined by changing the stimulating intensity from 1.5 to 6 V. The initial slope of the EPSPs was measured. Field EPSPs arising from the MF synapses were evoked by stimulating the dentate granule cell layer and were recorded from the stratum lucidum of CA3. The amplitude of field EPSPs was measured with a 0.5-ms window positioned at 70–80% of the peak of the baseline field EPSPs. More than 85% block of EPSP by an agonist of group II metabotropic glutamate receptors (2*S*,2’*R*,3’*R*)-2-(2’,3’-dicarboxycyclopropyl) glycine (DCG-IV, 1 μM) was the criterion used to identify MF input. A single electrical stimulation was delivered at a frequency of 0.05 Hz. All recordings were performed using a Multiclamp 700 B amplifier (Molecular Devices, Sunnyvale, CA, USA), filtered at 2 kHz, and stored on a personal computer via an interface (digitized at 10 kHz). Data were obtained from distinct samples. Serotonin hydrochloride was purchased from Sigma-Aldrich.

### Sample isolation for immunohistochemistry

Five’-bromo-2’-deoxyuridine (BrdU, 150 mg/kg, Nacalai Tesque, Japan) was intraperitoneally administered to mice once 2 h before sacrifice to evaluate cell proliferation to label the dividing cells, or four times on 14 and 15 days (twice a day) before sacrifice to evaluate cell survival.

For Dsp immunostaining, the mice were deeply anesthetized and transcardially perfused with cold saline. The brains were then stored at −80°C until further use. Serial fresh frozen sections (18 µm thick) were cut through the entire hippocampus using a cryostat (Leica 1510, Leica Microsystems, Wetzlar, Germany) and stored at −80°C. For other immunostaining procedures, mice were deeply anesthetized and perfused with cold saline followed by 4% paraformaldehyde (Sigma-Aldrich) in 0.1 M phosphate buffer (pH 7.4) and were post-fixed at 4 °C for 48 h. The brains were then cryoprotected in 20% sucrose for 48 h and in 30% sucrose for 24 h and stored at −80°C until further use. Serial sections (30 µm thick) were cut through the entire hippocampus using a cryostat and stored in 30% glycerol and 30% ethylene glycol in 0.02 M phosphate buffer (pH 7.4) at −20 °C until staining.

### Immunohistochemistry

For Dsp/NeuN or Dsp/NeuroD1 immunostaining, fresh frozen sections were fixed with 100% methanol for 20 min at 4 °C. After fixation, the sections were washed with phosphate-buffered saline (PBS), blocked using 10% equine serum in PBS, and incubated with monoclonal mouse anti-Dsp (1:50; Progen, 61003, RRID:AB_2920666) and rabbit anti-NeuN (1:500, Abcam, ab177487, RRID:AB_2532109) or rabbit anti-NeuroD1 (1:500; Abcam, ab213725, RRID:AB_2801303) at 4 ℃ overnight. After washing with PBS, the sections were incubated with donkey anti-mouse IgG antibody conjugated to Alexa Fluor 488 (1:300; Thermo Fisher Scientific, RRID:AB_141607) and donkey anti-rabbit IgG antibody conjugated to Alexa Fluor 555 (1:1000; Thermo Fisher Scientific, RRID:AB_162543) for 60 min. The sections were then incubated with 4’,6-diamidino-2-phenylindole (DAPI; 1:5,000; Merck Millipore) and covered with Mowiol (Merck Millipore) after washing with PBS. Confocal images were taken using Olympus FV3000. Single confocal images were shown for localization of Dsp. Anti-Dsp and rabbit anti-mouse IgG antibody conjugated to Alexa 555 (1:300; Thermo Fisher Scientific, RRID:AB_2535848) were used to confirm Dsp knockdown.

For FosB, NeuroD1, SOX2, and doublecortin (DCX) immunostaining, the 30 µm thick floating sections were washed with PBS and blocked using 10% equine serum (SH30074; Cytiva) in PBS containing 0.3% Triton X-100 (PBST) at room temperature for 60 min, followed by overnight incubation with rabbit anti-FosB (1:1000; Abcam, ab184938, RRID: AB_2721123), goat anti-SOX2 (1:200; Santa Cruz, sc17320, RRID:AB_2286684), rabbit anti-NeuroD1 (1:1000), rabbit anti-doublecortin (DCX) (1:1000; Cell Signaling Technology, Danvers, MA, USA; RRID:AB_561007) at 4 °C. For BrdU or BrdU/NeuN immunostaining, the floating sections were incubated in 50% formamide in 2×saline sodium citrate buffer (SSC) for 2 h at 60 °C, followed by incubation in 2 M hydrogen chloride at 37 °C for 30 min, and neutralized with 0.1 M boric acid (pH 8.5) at room temperature for 10 min before blocking. After blocking, the sections were incubated overnight with monoclonal rat anti-BrdU (1:8000; Abcam, ab6326, RRID:AB_305426) and rabbit anti-NeuN (1:500) at 4°C. After washing with PBST, the sections were incubated with donkey anti-rabbit IgG antibody conjugated to Alexa Fluor 555 (1:1000 for NeuroD1 and FosB immunostaining; Thermo Fisher Scientific, RRID:AB_2535853) or conjugated to Alexa Fluor 647 (1:1000 for NeuN immunostaining; Thermo Fisher Scientific, RRID: RRID:AB_162542), donkey anti-goat IgG antibody conjugated to Alexa Fluor 555 (1:1000 for SOX2 immunostaining; Thermo Fisher Scientific, RRID:AB_141788) and donkey anti-rat IgG antibody conjugated to Alexa Fluor 555 (1:1000 for BrdU immunostaining; Abcam, RRID: AB_2813834) for 60 min. After washing with PBST, the sections were incubated with DAPI (1:10,000) and mounted on slides with Mowiol after washing with PBS. For DCX immunostaining, the sections were incubated with biotinylated goat anti-rabbit IgG (1:200; Vector Laboratories Inc., Burlingame, CA, USA; BA1000, RRID: AB_2313606) for 60 min. The sections were washed with PBST and incubated with the ABC Vectastain Kit (Vector), and antigen detection was performed with 0.06% 3,3’-diaminobenzidine (Wako) staining. After washing, the sections were mounted on slides using Entellan New (Merck Millipore). Sections were photographed under a microscope (BZX-700, Keyence, Osaka, Japan).

### Quantification of BrdU-, SOX2-, NeuroD1-, DCX-, and FosB-positive cells

Computer-assisted image analysis (ImageJ, NIH, Bethesda, MD, USA; QuPath, Bankhead et al., 2017) was used for cell analysis. For quantification of FosB-positive cells, three hippocampal sections with EGFP-expressing DG were photographed. The number of FosB-positive cells was counted and presented as the number of cells per 10,000 μm^2^ area in the granule cell layer of the DG. For BrdU-labeled cell quantification, 6–8 hippocampal sections with EGFP-expressing DG were photographed. To evaluate cell proliferation, the number of BrdU-positive cells was counted in the subgranular zone (SGZ) and presented as the number of cells per 1,000 μm along the SGZ. To evaluate cell survival, the number of BrdU-positive cells in the granule cell layer of the DG was counted and presented as the number of cells per 100,000 μm^2^ of the granule cell layer. To evaluate NeuN colocalization of BrdU (+) cells, z-sectioning was used to determine if BrdU (+) cells were colabeled with NeuN. At least 30 BrdU (+) cells per animals were analyzed for co-localization with NeuN. For SOX2- or NeuroD-positive cell quantification, 3–4 hippocampal sections with EGFP-expressing DG were photographed. The number of SOX2- or NeuroD1-positive cells in the SGZ was counted and presented as the number of cells per 1,000 μm along the SGZ. For DCX-positive cell quantification, 2–3 hippocampal sections with EGFP-expressing DG were photographed. DCX-positive cells in which dendrites reached the molecular layer and underwent complex processes were subcategorized as DCX-positive cells with long and branched dendrites. The total number of DCX-positive and DCX-positive cells with long dendrites was counted in the SGZ and presented as the number of cells per 100 μm along the SGZ.

### Golgi staining

Golgi staining was performed using a FD Rapid Golgi Stain Kit (PK401, FD Neuro Technologies) according to the manufacturer’s instructions. Briefly, mice were sacrificed by decapitation at 24 h after the social interaction test, and the brains were quickly collected in MilliQ water. The hippocampal part of the brain was dissected and immersed in freshly prepared Golgi-Cox impregnation buffer included in the kit. The impregnation buffer was refreshed after 24 h, and the brains were incubated for 2 weeks. After cryoprotection with the cryoprotectant buffer included in the kit at 4°C for 1 week, each brain was frozen and kept at 80°C and cut into 100-μm sections with a cryostat. The sections were mounted on gelatin-coated slides, dried for 72 h, and stained according to the manufacturer’s instructions. The slides were dehydrated in ethanol, cleared in xylene, and covered with Entellan New. The sections were photographed under a microscope (BZX-700, Keyence, Osaka, Japan). The total number of spines in dendrites of Golgi-impregnated granule neurons in the inner molecular layer in the DG were counted. ImageJ was used to measure dendritic length. Analysis was performed on 10 dendrites in per mouse (n=6 each group).

### Behavioral experiments

The mice were habituated to the testing room for at least 30 min before the behavioral test. During the test, the movement of each mouse was recorded using a digital video camera. All records were stored and analyzed using the SMART 3.0 video tracking software (Panlab, Barcelona, Spain, RRID: SCR_002852). The apparatus was thoroughly cleaned with 70% ethanol after the removal of each mouse.

The open field test (OFT) was performed for 10 min using an apparatus composed of opaque white walls and a floor (50 cm × 50 cm × 20 cm) illuminated at an intensity of 80 lux in the center of the floor. Each mouse was placed in the center of an open-field arena, and locomotor activity was recorded for 10 min. The total distance moved, relative time spent in the central zone, and entry time into the central zone were measured. The floor was divided into 25 squares, and the four central squares were defined as the central zone.

The forced swim test (FST) was performed in a cylindrical container (13 cm diameter, 25.5 cm height) filled with water to a height of 18 cm. Water (23–25 °C) was replaced between the trials. The FST was performed and recorded for 10 min using a digital video camera. The duration of immobility was then quantified.

The spontaneous locomotor activity (LMA) test was performed in a transparent home cage without bedding (20 cm length × 12.5 cm width) over 60 min. The total distance moved during the last 30 min was tracked.

The compartment social interaction test (SIT) was performed in an SIT apparatus (LE894T, Panlab) consisting of three chambers (42 cm × 60 cm × 22 cm) and two partitions (5.5 cm × 0.15 cm × 17 cm). Each mouse was placed in the middle chamber and allowed to explore the chamber freely for 5 min. Immediately after habituation, an unfamiliar mouse (a 7–8-week-old male C57BL/6N mouse) was introduced into one of the two side chambers enclosed in a wire cage. An identical empty wire cage was placed on the opposite side of the chamber. After placement, the side doors were opened simultaneously, and the test mouse was allowed to explore the entire three-chamber arena for 10 min. The time spent in each compartment was measured.

An elevated plus maze (EPM) test was performed in an EPM apparatus (EPM-X, Muromachi Kikai, Japan) consisting of two open arms (30 cm × 6 cm, 40 lux) and two closed arms (30 cm × 6 cm × 15 cm, 20 lux) extending from a central platform (6 cm × 6 cm). The maze was elevated to 40 cm above the floor. Each mouse was placed in the center of the maze, facing the open arm, and allowed to explore freely for 10 min. The total distance and time spent in the open arms were measured.

## Statistical analyses

All data are presented as the mean ± standard error of the mean. Statistical analyses were performed using an unpaired Student’s *t*-test, Welch’s *t*-test or two-way analysis of variance (ANOVA). Statistical significance was set at *P* < 0.05. Before performing *t*-test, normality of the data was examined using the Shapiro–Wilk test. Detailed statistical data are presented in Table 2. Outliers in data from Fig. 5B were removed using the Grubb’s outlier test. Otherwise, we did not exclude samples from statistical analysis. All analyses were performed using the PRISM 9 software (GraphPad, San Diego, CA, USA).

**Table 2.**
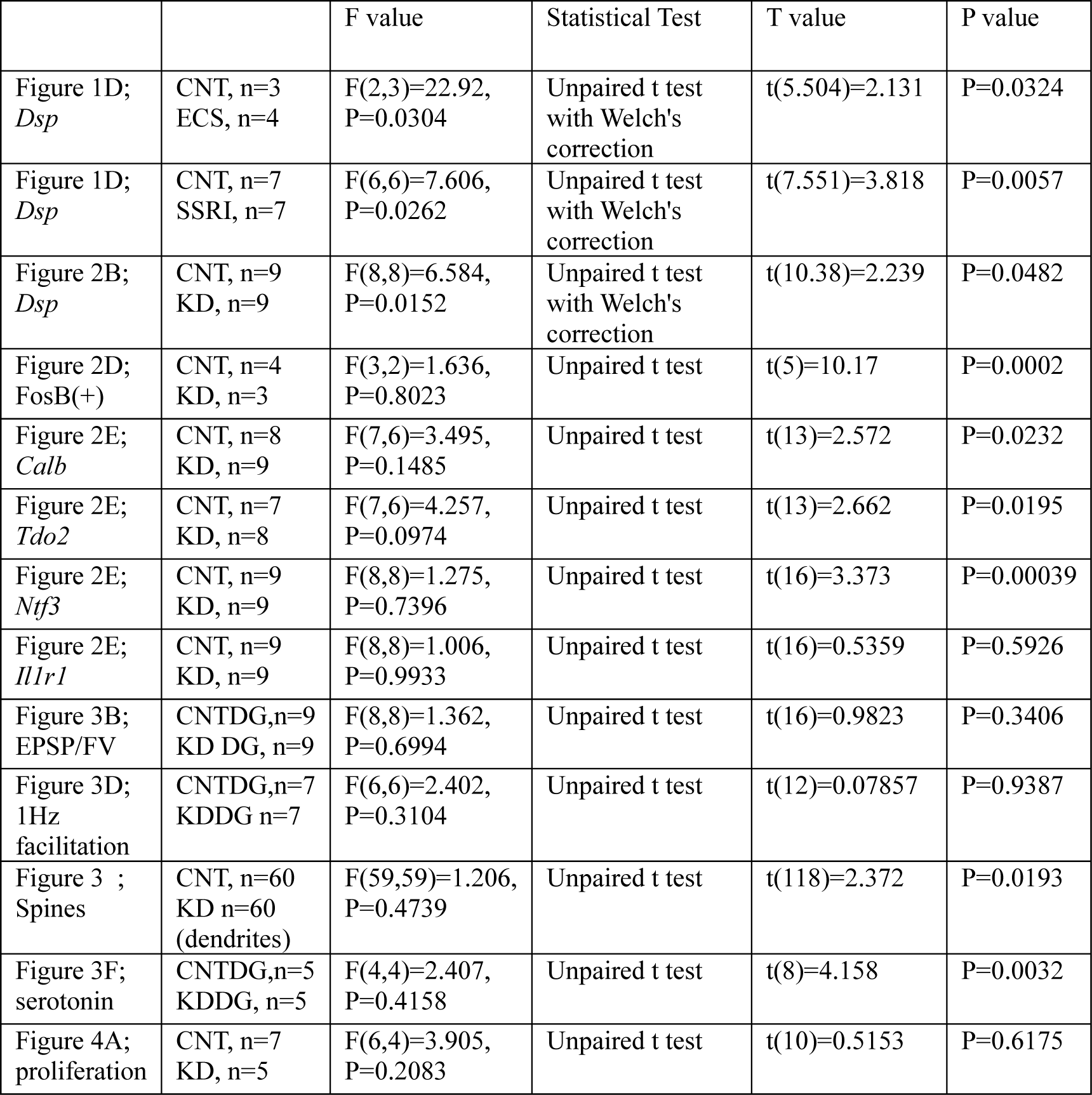
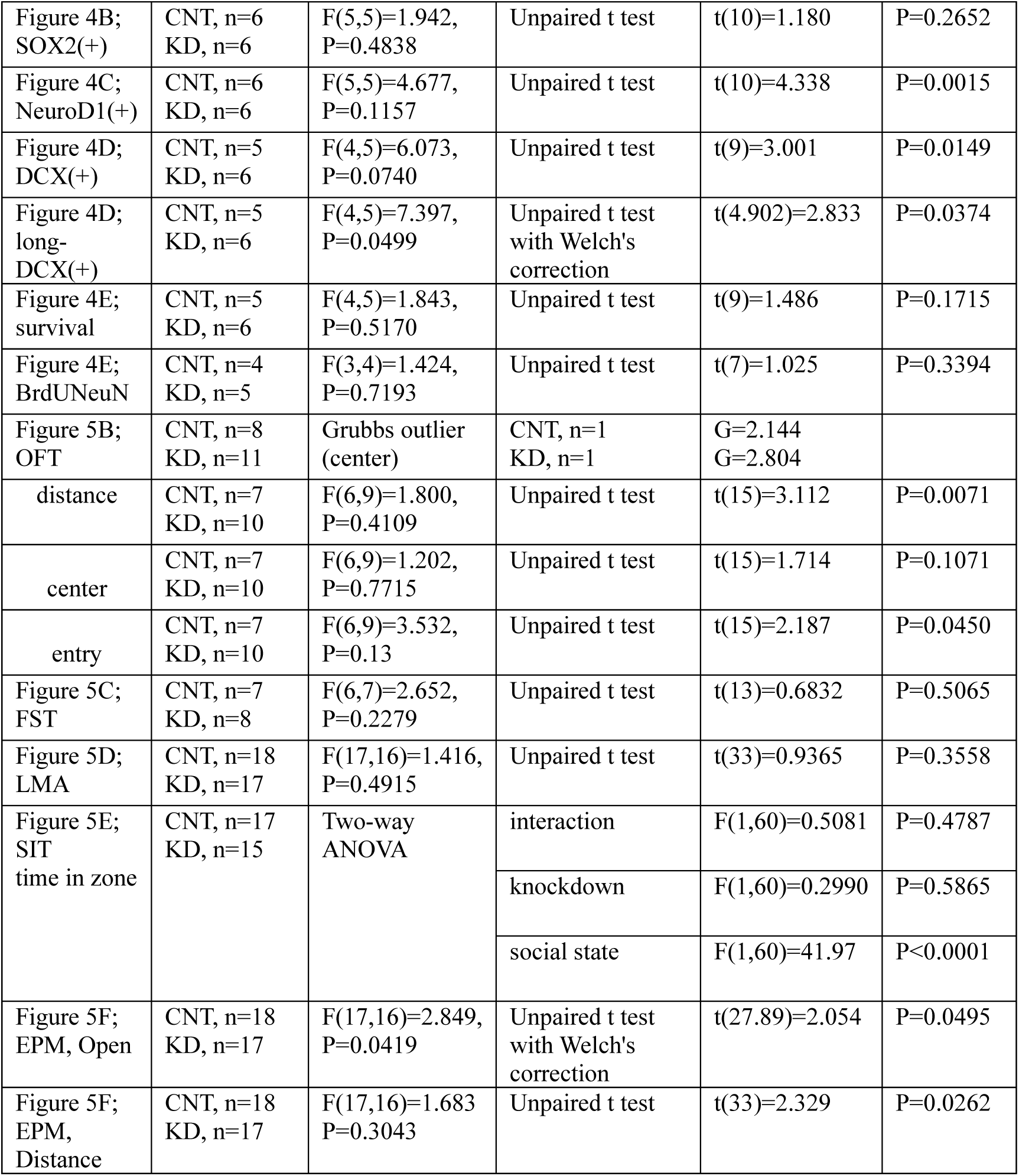
Statistical analysis.

## Results

### Expression of Dsp in the hippocampal DG and its regulation by antidepressant stimulation

*Dsp,* a granule cell-specific maturation marker (Yamasaki et al., 2008; Kobayashi et al., 2010), is highly and specifically expressed in the DG of the hippocampus of the central nervous system (Fig. 1A, Allen Institute for Brain Science, 2004). However, the localization of Dsp protein in the DG has not been fully investigated. Therefore, we initially examined the cellular localization and cell type of Dsp expression. Immunohistological analysis revealed that Dsp was highly expressed in the granule cell layer and molecular layer in the DG compared to the CA1 region (Fig. 1B). Dsp immunoreactivity signals were detected in NeuN-positive cells in the DG (Fig. 1C, upper). However, Dsp signals were rarely detected in NeuroD1-positive neural progenitors or immature neurons in the SGZ of the DG (Fig. 1C, lower, arrows). These findings suggest that Dsp is expressed in mature granule neurons in the DG. Dsp immunostaining was only observed after methanol fixation; therefore, the localization of other neuronal markers such as an immature neuronal marker DCX was not investigated.

**Figure 1.**
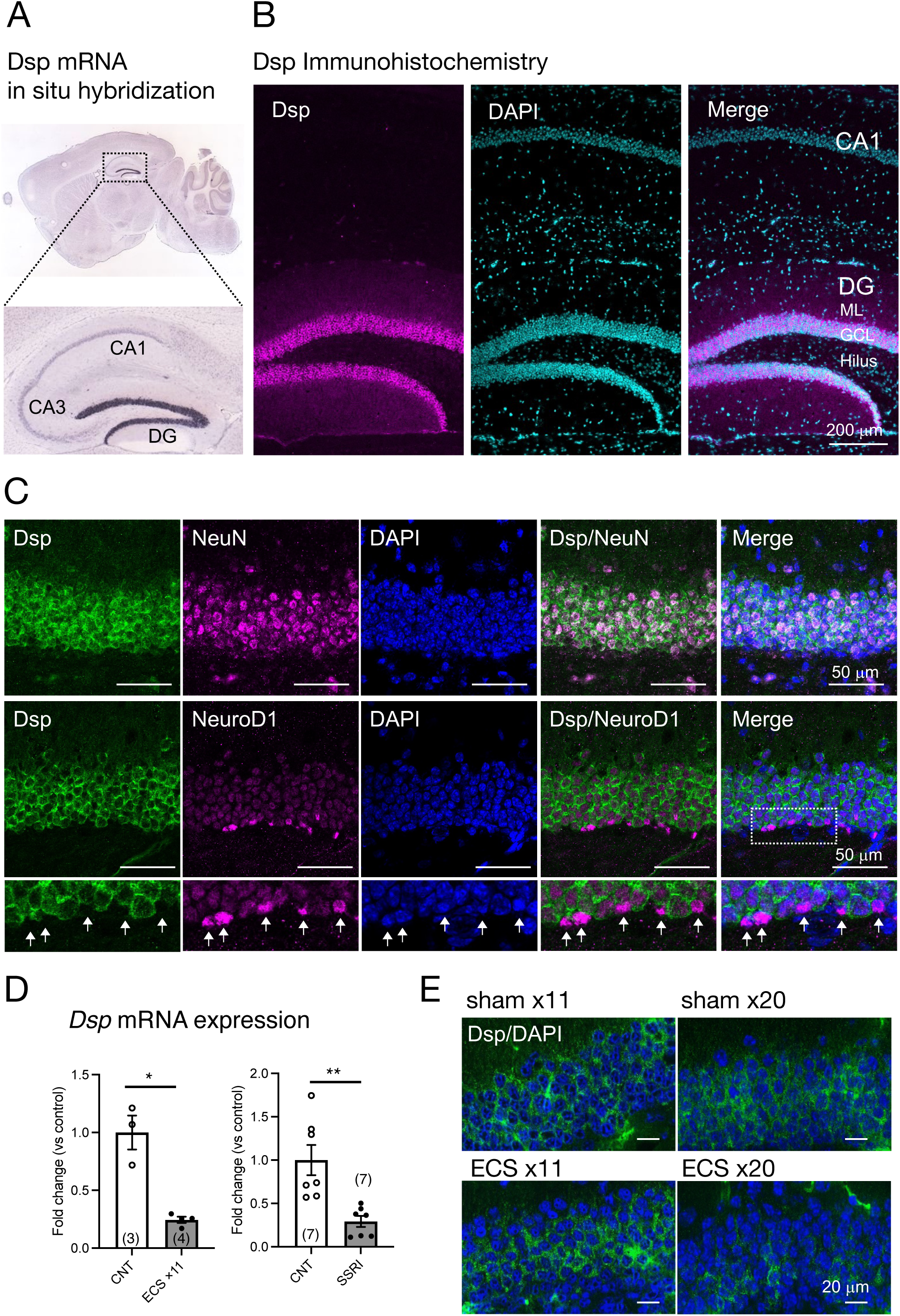
Expression of Dsp in the dentate gyrus (DG) of the hippocampus. (A) Expression of Dsp mRNA in adult mouse brain from Allen Mouse Brain Atlas. (B) Low magnification images of immunoreactivity for Dsp (magenta) in the DG and CA1. DAPI (cyan) is utilized for nuclear staining. GCL; granule cell layer, ML; molecular layer. Scale bar: 200 μm. (C) Representative images of immunoreactivity for Dsp (green), NeuN (upper, magenta), and NeuroD1 (lower, magenta) in the DG. DAPI was used for nuclear staining (blue). Enlarged images of the dotted square are shown below. Arrows represent Dsp-negative and NeuroD1-positive cells (arrows). Scale bars: 50 μm. (D) Relative gene expression of *Dsp* at 24 h after 11 times of electroconvulsive seizure (ECS) stimulation (left, P=0.0324) or 4-weeks treatment of selective serotonin reuptake inhibitor (SSRI) fluoxetine (right, P=0.0057). Data are expressed using dot plots and means ± standard error of mean. The n number is given in graph. *, *P*<0.05; **, *P*<0.01. Detailed statistical data are shown in Table 2. (E) Representative images of immunoreactivity for Dsp (green) in the DG after 11 or 20 times of ECS stimulation. Scale bars: 20 μm.

Furthermore, we investigated whether Dsp expression was regulated by antidepressant stimulation at the protein level. The expression of *Dsp* in the DG was reduced by chronic administration of electroconvulsive seizure (ECS) or a selective serotonin reuptake inhibitor (SSRI) (Fig. 1D), as previously reported (Imoto et al., 2017). Immunostaining analysis showed that Dsp protein began to decrease in outer and inner granule neurons at 11 times of ECS, and its expression almost disappeared at 20 times of ECS administration (Fig. 1E). Thus, suppression of desmoplakin protein expression by antidepressant treatments would be expected to require a longer time or stimulus than suppression of mRNA expression.

### Dsp expression knockdown in the DG by an adeno-associated virus (AAV) carrying Dsp miRNA

To explore the role of Dsp in hippocampal function, we performed an AAV-mediated knockdown of Dsp in the DG. For this purpose, we generated AAVrh10 expressing EGFP and an artificial miRNA targeting Dsp mRNA (AAV-Dsp miRNA; Fig. 2A). The AAV rh10 serotype shows high transduction efficiency into granule neurons in the DG (Sehara et al., 2022). Seven to eight weeks after the injection of AAV-Dsp or AAV-control miRNA into the DG, strong EGFP signals, which were co-expressed with miRNA, were observed in the molecular layer and granule cell layer in the DG and CA3 projection area (Fig. 2A, left). EGFP signals did not co-localize in DCX-positive cells in the DG (Fig. 2A, right), indicating that AAV-mediated gene transfer occurs primarily in mature granule neurons of the DG. *Dsp* mRNA expression significantly decreased in the AAV-Dsp miRNA group (Fig. 2B). Moreover, immunohistological analysis showed a decrease in Dsp immunoreactivity in the DG following AAV-Dsp miRNA injection (Fig. 2C), demonstrating that the sequence of artificial miRNAs targeting Dsp used in this study has the potential to knockdown Dsp expression.

**Figure 2.**
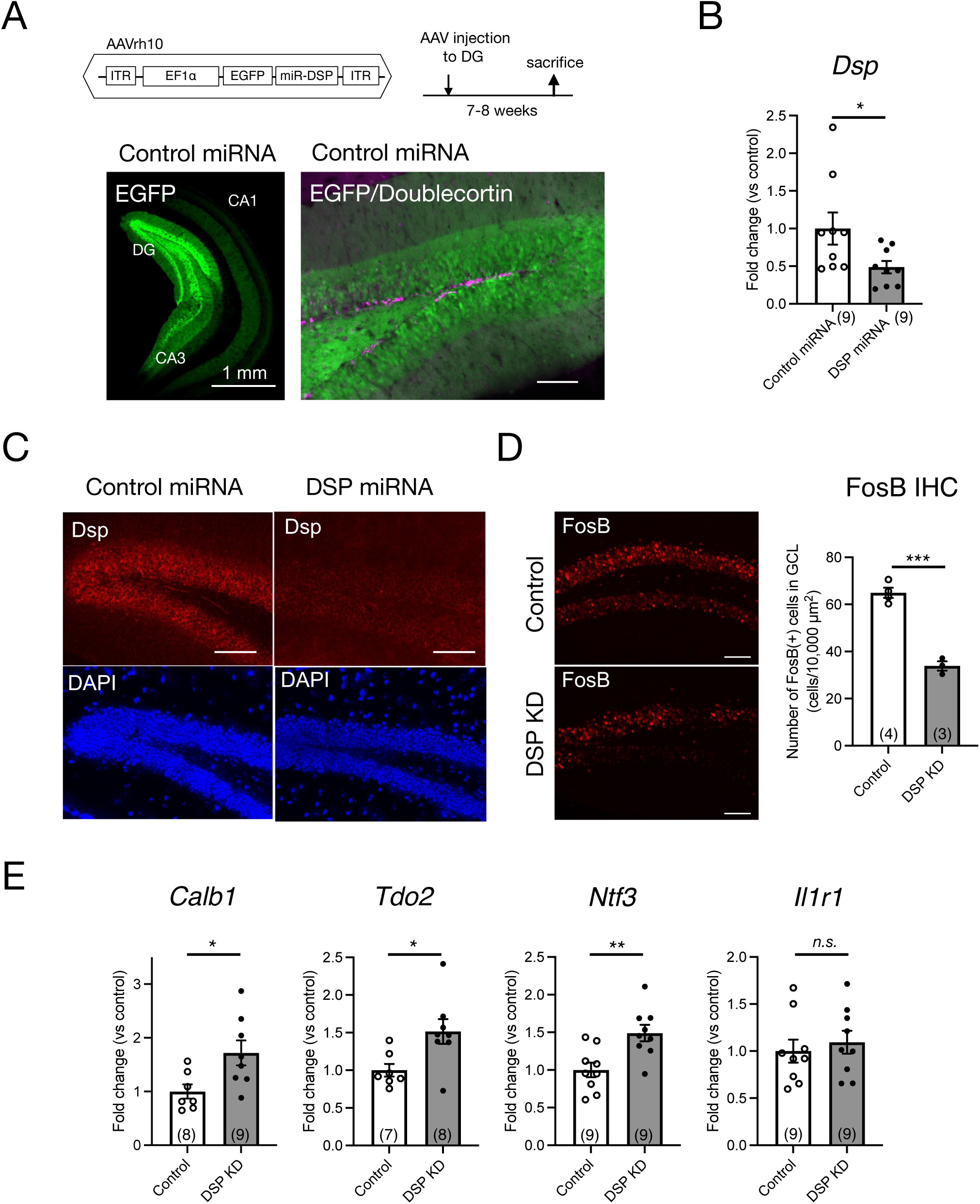
Knockdown of Dsp in the DG by adeno-associated virus (AAV) carrying Dsp artificial miRNA and expression changes in the DG. (A) Structure of AAVrh10 carrying EGFP and artificial microRNA sequence for Dsp (DSP miRNA) and time course of experimental procedure. Representative EGFP image 7–8 weeks after control miRNA AAV injection into the DG (left). EGFP and immunoreactivity for an immature neuronal marker doublecortin (magenta) in the DG (right). (B) Gene expression of *Dsp* 7–8 weeks after AAV infection (P=0.0482). (C) Representative images of immunoreactivity for Dsp (red) and DAPI (blue) in the DG after AAV infection. (D) Effect of Dsp knockdown in the DG on the expression of FosB protein. Representative images of immunoreactivity for FosB in the DG (left). Quantification of FosB-positive cell numbers (right) in the granule cell layer (P=0.0002). (E) Effect of Dsp knockdown in the DG on the expression of DG specific maturation markers in the DG. Expression of *Calb1* (P=0.0232)*, Tdo2* (P=0.0195)*, Ntf3* (P=0.00039), and *Il1r1* (P=0.5926). Data are expressed using dot plots and means ± standard error of mean. The n number is given in graph. *, *P*<0.05; **, *P*<0.01; ***, *P*<0.001; n.s., not significant. Detailed statistical data are shown in Table 2. Scale bars: 1 mm for the low magnification image in Fig. 2A (left); 100 μm for other images.

### Changes in activity marker expression and spine number in the DG and alteration of the sensitivity to serotonin at the MF synapse by Dsp knockdown

First, we examined the direct effect of Dsp knockdown on the activity of dentate granule neurons. FosB, including ΔFosB, is induced in response to many types of chronic stimuli and accumulates because of its high stability (Nestler et al., 1999; Lamothe-Molina et al., 2022). Consequently, we performed immunohistochemical analysis using an anti-FosB antibody to monitor neuronal activity in the DG under normal home-cage conditions. FosB immunoreactivity in the granule cell layer of the DG significantly decreased in the Dsp knockdown group (Fig. 2D).

Because Dsp is a granule cell-specific DG maturation marker (Yamasaki et al., 2008; Kobayashi et al., 2010), we examined the expression of other markers, including *Calb, Tdo2, Ntf3, and Il1r1* in the DG. The expression of *Calb, Tdo2,* and *Ntf3* significantly increased in the DG of the Dsp knockdown group (Fig. 2E).

Electrophysiological analyses of synaptic transmission were further performed using acute hippocampal slices (Fig. 3A-I). We first examined the input-output relationship of MPP-granule cell synaptic transmission by changing the intensity of stimulation (Fig. 3A). No significant change in the relationship between fiber volley amplitude and EPSP slope was observed (Fig. 3B and C). Next, properties of DG output MF-CA3 synaptic transmission were examined (Fig. 3D) and there was no change in the ratio of field EPSP to fiber volleys (Fig. 3E). These results suggest that the basal transmission efficacy at the MPP-granule cell and the MF-CA3 synapses was not altered by Dsp knockdown. Antidepressant treatment attenuates the synaptic facilitation induced by repetitive stimulation and augments serotonergic modulation at the MF synapse (Kobayashi et al., 2010; Imoto et al., 2017; Kobayashi et al., 2020). Dsp knockdown did not change synaptic facilitation after 1 Hz stimulation (Fig. 3F and G). Bath-applied serotonin induced the potentiation of field EPSPs in the AAV control group (Fig. 3H), consistent with the findings of previous studies. The magnitude of serotonin-induced potentiation was significantly reduced in the Dsp knockdown group (Fig. 3H and I). To further investigate the role of Dsp in spine formation in the granule neurons, we employed Golgi staining to assess spine number of granule cell dendrites in the inner molecular layer (Fig. 3J). Dsp knockdown slightly but significantly decreased the total number of spines (Fig. 3K).

**Figure 3.**
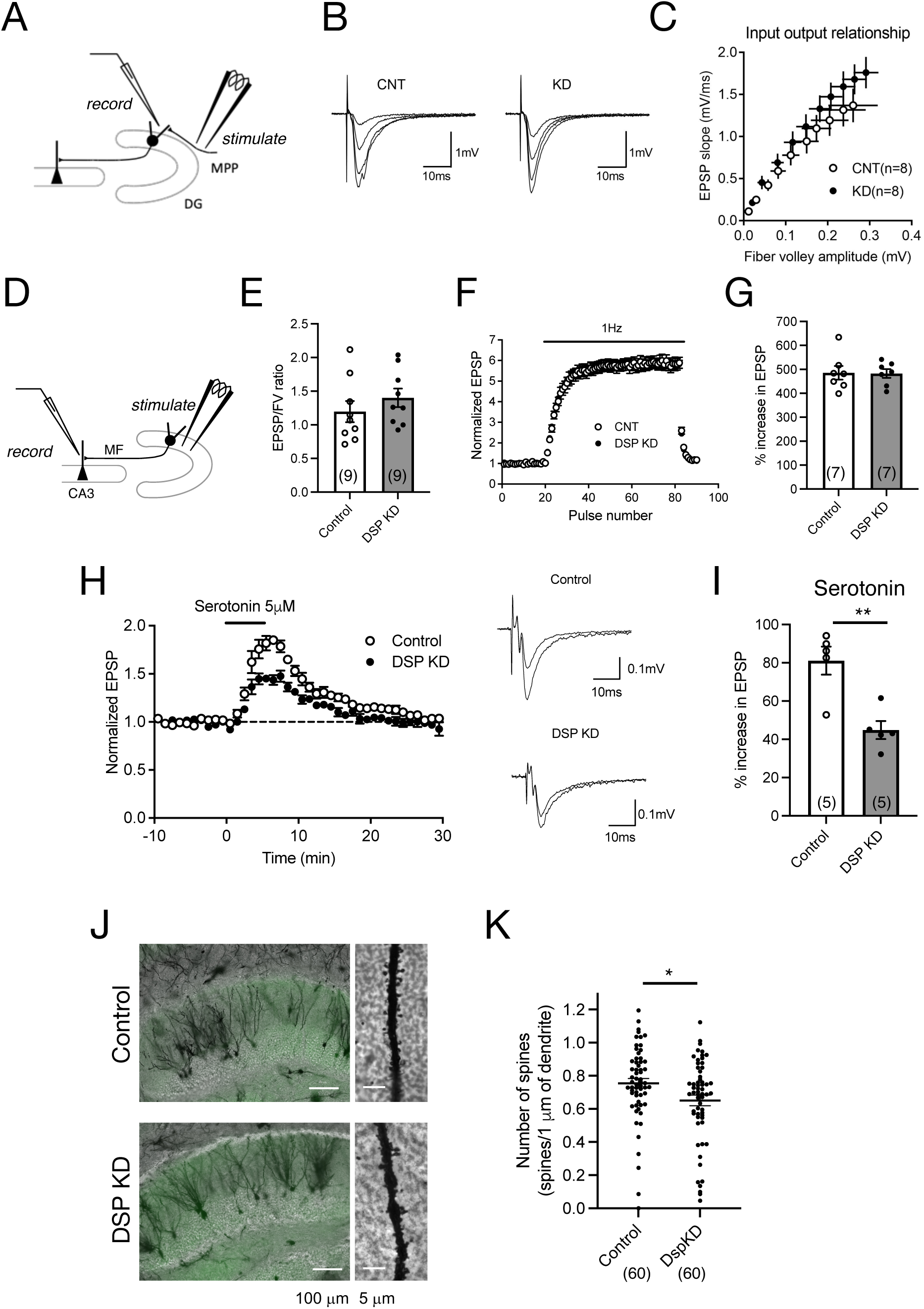
Electrophysiological changes in the DG by Dsp knockdown. (A) Diagram showing the electrode arrangement for recording field excitatory post synaptic potentials (EPSPs) at the medial perforant path (MPP)–granule cell synapse. (B) Sample recordings show MPP synaptic potentials evoked at the stimulus intensities of 2, 3, 4 and 5 V in control DG and knockdown DG. (C) The relationship between fiber volley and EPSP at MPP–DG synapse. (D) Diagram showing for recording field EPSPs at the mossy fiber (MF)–CA3 synapse. (E) Ratios for MF field EPSPs to fiber volley (FV) at the baseline stimulus intensity (P=0.3406). (F and G) Effect of Dsp knockdown on synaptic facilitation induced by repetitive stimulation at the MF synapse (G, P=0.9387). (H) Effect of Dsp knockdown on potentiation of MF synaptic transmission induced by serotonin (5 μM, 5 min) applied in the bath. Sample recordings show averages of nine consecutive field EPSPs before and at the end of serotonin application in control DG and knockdown DG. (I) Effect of Dsp knockdown on serotonin-induced synaptic potentiation (P=0.0032). (J) Representative images of Golgi staining in the DG (left) and dendrites (right) in inner molecular layer. (K) Quantitative analysis of the number of spines in the dendrites. Ten dendrites per animal and six mice per group were analyzed. Data are expressed using dot plots and means ± standard error of mean. The n number of data is given in graph and represents the number of slices (E, G, I) and dendrites (K). **, *P*<0.01. Detailed statistical data are shown in Table 2.

### Changes in neurogenic activity in the DG by Dsp knockdown

Adult hippocampal neurogenesis consists of proliferation, differentiation, and survival and is regulated by various factors, including signals from adjacent mature granule neurons in the DG (Kempermann et al., 2015). Since the above results suggest that Dsp regulates neuronal activation in the DG, we explored the influence of Dsp knockdown on hippocampal neurogenesis. First, we examined the effect of Dsp knockdown on cell proliferation. BrdU was administered 2 h before the sacrifice to label proliferating cells (Fig. 4A, left). BrdU-positive cells were detected in the SGZ of the DG (Fig. 4A, middle), and no difference was observed in the number of proliferating cells between the control and Dsp knockdown groups (Fig. 4A, right). We next examined the number of SOX2-positve cells in the SGZ. SOX2 is a marker for neural stem cells and progenitors in the adult DG (Segi-Nishida et al., 2008). There was no difference in the number of SOX2-positive cells in the SGZ (Fig. 4B). We then examined the expression of Neuro D1, which is expressed in progenitors/neuroblasts and immature neurons in the adult DG. The number of NeuroD1-positive cells in the SGZ was significantly decreased in the Dsp knockdown group (Fig. 4C). We further examined the expression of the immature neuronal marker, DCX, in the DG. The total number of DCX-positive cells and those with long dendrites decreased in the Dsp knockdown group (Fig. 4D). Because most newborn cells in the DG die within 2–3 weeks following proliferation in mice (Ueno et al., 2019), we evaluated the effect of Dsp knockdown on the survival of newborn cells 2 weeks after BrdU administration (Fig. 4E, left). The number of BrdU-positive cells in the DG showed a tendency to decrease, yet there was no significant difference between the control group and the Dsp knockdown group (Fig. 4E, middle). Furthermore, there was no difference in the ratio of NeuN-positive cells among the BrdU-positive cells.

**Figure 4.**
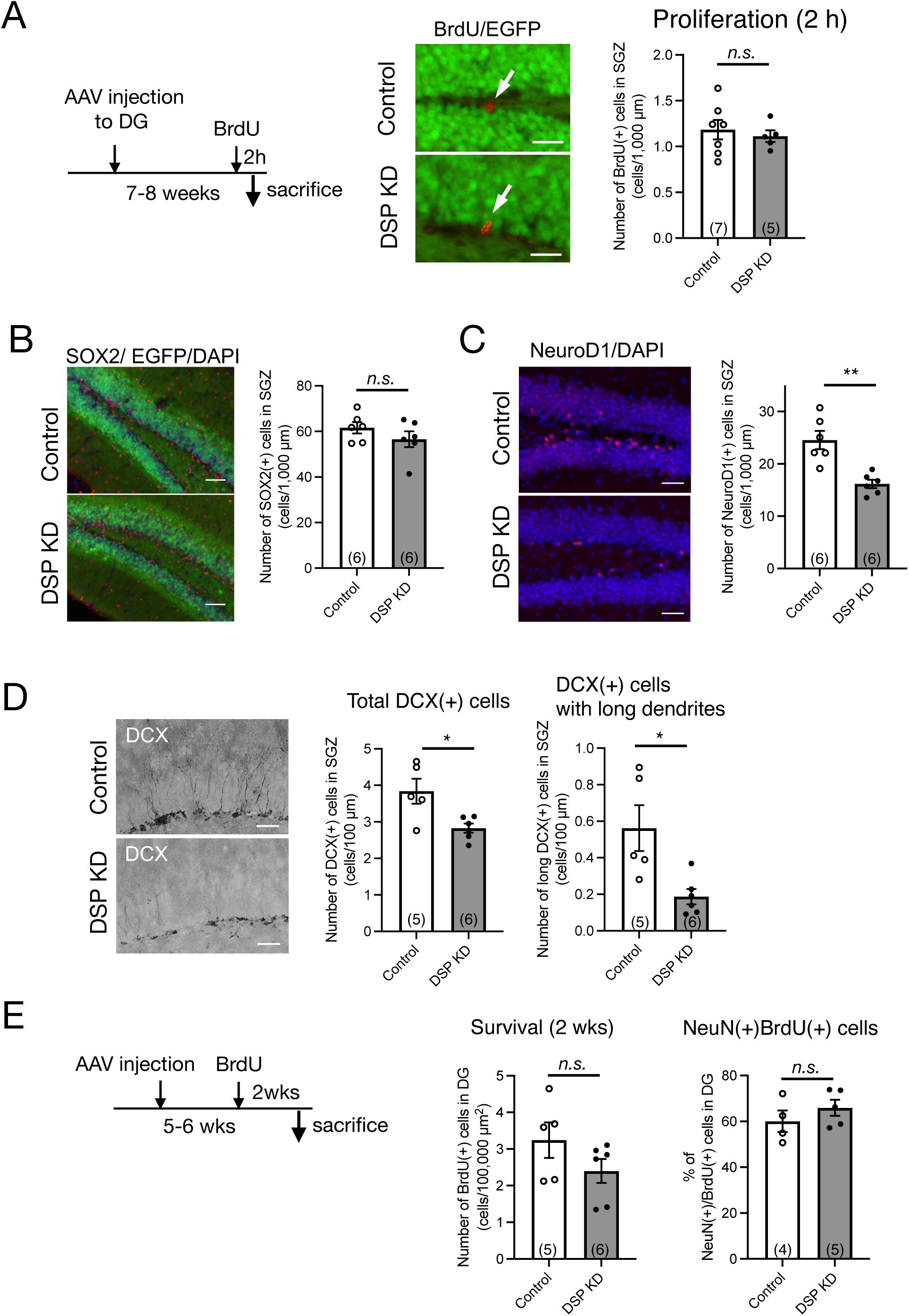
Adult neurogenic changes in the DG by Dsp knockdown. (A) Effect of Dsp knockdown on proliferation in the DG. Left: Time course of the experimental procedure. Middle: Representative images of immunoreactivity for BrdU (red). The arrows represent BrdU-positive cells in the SGZ. Scale bars: 20 μm. Right: Quantification of BrdU-positive cell numbers in the SGZ (P=0.6175). (B) Effect of Dsp knockdown on the number of neural stem cells or progenitors in the DG. Left: Representative images of immunoreactivity for SOX2 in the DG. Scale bars: 100 μm. Right: Quantification of SOX2-positive cell numbers in the SGZ (P=0.2652). (B) Effect of Dsp knockdown on neuronal differentiation in the DG. Left: Representative images of immunoreactivity for NeuroD1 in the DG. Scale bars: 100 μm. Right: Quantification of NeuroD1-positive cell numbers in the SGZ (P=0.0015). (C) Effect of Dsp knockdown on cell number of doublecortin (DCX)-positive immature neurons in the DG. Left: Representative images of immunoreactivity for DCX in the DG. Scale bars: 40 μm. Middle: Quantification of DCX-positive cell numbers in the SGZ (P=0.0149). Right: Quantification of DCX-positive cells with long dendrites in the SGZ (P=0.0374). (E) Effect of Dsp knockdown on survival of 2-weeks of cell age in the DG. Left: Time course of the experimental procedure. Middle: Quantification of BrdU-positive cells of 2-weeks (2 wks) of cell age in the DG (P=0.1715). Right: The ratio of NeuN(+) cells among the BrdU (+) cells in the DG (P=0.3394). Data are expressed using dot plots and means ± standard error of mean. The n number is given in graph. *, *P*<0.05; **, *P*<0.01. n.s. not significant. Detailed statistical data are shown in Table 2.

**Figure 5.**
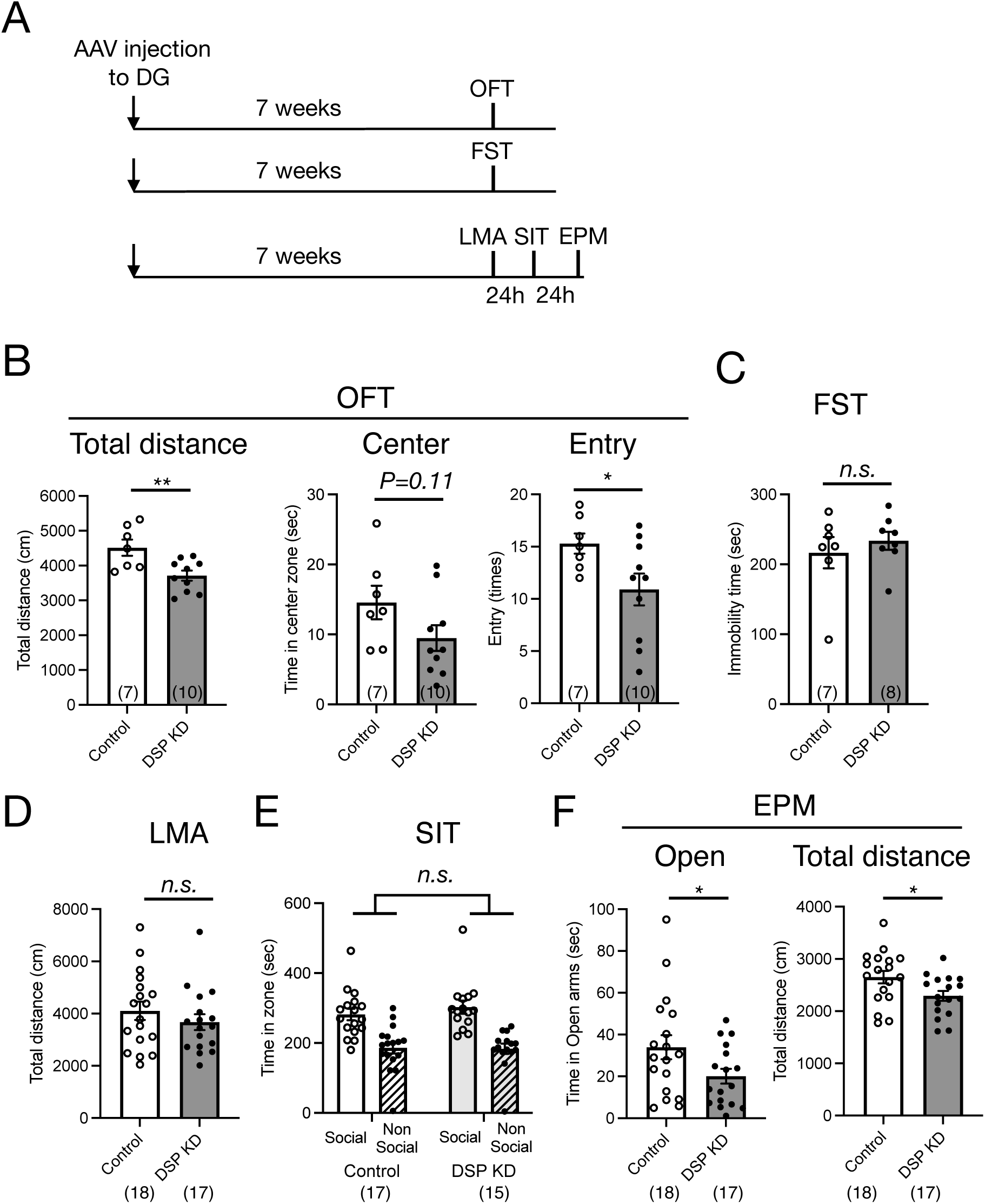
Behavioral changes in the DG by Dsp knockdown. (A) Time course of the experimental procedure. (B) Exploratory behaviors of open field test (OFT). Total distance (left, P=0.0071) in the open field. Time spent (middle, P=0.1071) and entry times (right, P=0.0450) in the center of the open field. (C) Immobility time in the forced swim test (FST, P=0.5065). (D) Total distance of locomotor activity (LMA) in the home cage (P=0.3558). (E) Social interaction behavior in the 3-chamber social interaction test (SIT). A stranger mouse within a wire cup placed in one chamber (social) and an empty wire cup placed in another chamber (Non-Social). (F) Time spent on the open arms (left, P=0.0495) and total distance (right, P=0.0262) in the elevate plus maze (EPM) test. Data are expressed using dot plots and means ± standard error of mean. The n number is given in graph. *, *P*<0.05; **, *P*<0.01; n.s. not significant. Detailed statistical data are shown in Table 2.

### Influence of Dsp knockdown on anxiety-related behaviors

The ventral DG is a key brain region involved in emotional behaviors, including anxiety- and depression-like behaviors (Sahay and Hen, 2007; Levone et al. 2015; Anacker et al. 2018). To examine the influence of Dsp knockdown in the DG on emotion-related behaviors, we conducted three sets of behavioral experiments (Fig. 5A). First, we performed an open-field test to assess behavior in an anxiety-provoking environment. Dsp knockdown in the DG suppressed locomotor activity and the number of entries into the central area in the open field box, and a suppressive trend was observed for the time spent in the central area (Fig. 5B). Next, we evaluated immobility time in the FST to assess depression-like behavior. No difference in immobility time between the control and Dsp knockdown groups was observed (Fig. 5C). Another set of experiments was conducted on locomotor activity in a home cage, social interaction test, and elevated plus maze test. Contrary to the data from the open field test, both control and knockdown groups demonstrated comparable locomotor activity under standard conditions (Fig. 5D), suggesting that the knockdown does not perturb baseline locomotor function. In the three-chamber social interaction test, the control mice spent more time around the area with an unfamiliar mouse than around the area with empty cages. This social interaction behavior was comparable to that observed in the Dsp knockdown mice (Fig. 5E). In the elevated plus maze test, a reduction in both the time spent in the open arm and locomotor activity was observed in Dsp knockdown mice (Fig. 5F). These results demonstrate that Dsp knockdown mice in the DG shows anxiety-like behavior. Also, these findings can implicate that expression of Dsp in the DG plays an anti-anxiety role.

## Discussion

This study aimed to investigate the role of Dsp expressed in the DG in hippocampal function and emotion-related behaviors. We showed that Dsp is localized mostly in mature granule neurons in the DG and that knockdown of Dsp decreases the expression of neuronal activity marker and serotonin responsiveness in DG neurons while enhancing the expression of mature neuronal markers. In addition, we demonstrated that Dsp knockdown in the DG alters adult neurogenic processes in the adjacent SGZ and that these hippocampal changes contribute to anxiety-like behaviors. These findings suggest that Dsp in the DG maintains the activity of mature neurons, regulates neuronal differentiation in the adult neurogenic process, and contributes to anxiolytic-like effects. This study is the first to demonstrate the role of Dsp in neuronal activity and emotion-related behaviors in the DG.

### Expression of desmoplakin in the DG

Since Dsp could play an important role in cell-to-cell adhesion and subsequent intracellular regulatory signaling, we initially confirmed its localization in the brain. Immunohistological studies showed that Dsp protein was localized in granule cell layer in the DG (Fig. 1B). Dsp colocalized with the NeuN-positive cells, but not with the neural progenitor/immature neuron marker NeuroD1-positive cells in the DG (Fig. 1C). These results are consistent with those of a previous study that demonstrated that Dsp colocalizes with the post-mitotic neuron marker Bcl11b but not with the neural stem cell/progenitor marker SOX2 in the DG at 14 days of postnatal age (Simon et al., 2012). Desmosome components include transmembrane cadherins and intracellular proteins such as plakoglobin, plakophilin, and Dsp (Garrod and Chidgrey, 2008; Broussard et al., 2015). Nevertheless, electron microscopy images of the rat DG showed no desmosome structures between the cell bodies (Laatsch and Cowan, 1966). Therefore, Dsp in the DG may form a complex different from that in epithelial desmosomes. The expression and interactions of each desmosome component in the DG will be examined to elucidate the localization and function of Dsp in neurons.

### Role of desmoplakin in mature granule neurons in the DG

We found that Dsp knockdown in the DG suppressed FosB expression (Fig. 2D); FosB is useful for monitoring steady-state activation. FosB, including ΔFosB, is induced in response to many types of stimuli and shows high stability (Nestler et al., 1999; Lamothe-Molina et al., 2022). In contrast, Dsp knockdown increased the expression of the mature neuronal markers *Calb1*, *Tdo2*, and *Ntf3* (Fig. 2E), which are negatively regulated by strong neuronal activation, such as in the ECS and epileptic seizures (Imoto et al., 2017; You et al., 2017). Although the number of dendritic spines in DG neurons was slightly but significantly reduced by Dsp knockdown, electrophysiological analysis showed that Dsp suppression did not alter the basal transmission efficiency at the MPP-granule cells. Excitatory inputs other than the MPP input may be altered, resulting in attenuated granule cell activity. It is also possible that Dsp knockdown affected inhibitory synapses or intrinsic excitability of granule cells.

Dsp suppression did not alter the basal transmission efficiency at the MF-CA3 synapses; however, it reduced serotonin-induced potentiation (Fig. 3). Synaptic potentiation by serotonin is mediated by presynaptic 5-HT4 receptor-cAMP signaling in the MF, which is enhanced by antidepressant treatment including ECS (Kobayashi et al., 2008; 2020). Dsp knockdown in the DG may reduce serotonin-mediated intracellular signaling in the DG.

Dsp is involved in various biological processes such as gene expression, differentiation, and cell migration (Bendrick et al., 2019; Dubash et al., 2016; Lombardi et al., 2016; Patel et al., 2014). Dsp interacts with plakoglobin, also known as γ-catenin (Delva et al., 2009). Suppression of Dsp expression in atrial myocyte cell lines leads to nuclear localization of plakoglobin/γ-catenin (Garcia-Gras et al., 2006). Further detail analysis will be required to characterize the Dsp-mediated transcriptional pathway. Especially, confirmation of the nuclear translocation of plakoglobin/γ-catenin will clarify whether changes in the expression of these factors are mediated by catenin signaling.

### Influence of desmoplakin knockdown on the neurogenic process in the DG

Adult neurogenesis in the SGZ of the DG is regulated by various signals from surrounding neurons (Kempermann et al., 2015). We investigated the role of Dsp in several neurogenic processes in the DG. Suppression of Dsp reduced the number of NeuroD1-positive neural progenitors and DCX-positive immature neurons, whereas the number of proliferating cells and SOX2-positive neural stem/progenitor cells remained unchanged (Fig. 4). EGFP signals, which should be coexpressed with miRNA, were not observed in DCX-positive cells of the DG (Fig. 2A), suggesting that Dsp in the DG indirectly regulates neurogenic processes in the adjacent SGZ. Since the rate of NeuN-positive neuronal differentiation at 14 days of cell age did not change between the control and Dsp knockdown, the period of NeuroD1 and DCX expression may be shortened. Thus, it is possible that Dsp knockdown promoted the rate of differentiation of newborn neurons, which allowed them to progress to maturity without sufficient differentiation. Forebrain neuron-specific ablation of Dsp impairs developmental hippocampal neurogenesis (Simon et al., 2012). During postnatal development in this mutant line, cell proliferation and neural differentiation in the DG were suppressed. All these findings suggest that Dsp is important for the regulation of cell proliferation during developmental neurogenesis and for neural differentiation during both developmental and adult neurogenesis.

How does Dsp indirectly regulate neuronal differentiation in post-mitotic neurons? Neuronal activation, such as that in the ECS, promotes the expression of adult neurogenic factors such as *Vegf*, *Fgf2*, and *Wnt2* in the DG (Newton et al., 2003; Warner-Schmidt et al., 2007). Conversely, the suppression of neuronal activity in the DG by inhibiting Dsp expression may reduce the expression of these growth factors. Furthermore, intracellular skeletal regulation by Dsp may also affect neurogenic niche formation. The neurogenic niche provides a unique milieu containing an extracellular matrix, humoral factors, and cell-cell contacts that allow controlled neuronal development (Kempermann et al., 2015). Altering the cytoskeleton of dentate granule neurons may affect intercellular contacts and extracellular matrix adhesion and inhibit neurogenic niche formation.

### Desmoplakin in the DG as a key regulator for emotion-related behaviors

The DG of the hippocampus controls emotion-related behaviors (Segi-Nishida and Suzuki, 2022). Although Dsp knockdown in the DG did not affect baseline locomotor function, depression-like behavior, or social behavior, it reduced the time spent in the open arm in the EPM test and reduced activity in the novel open field and EPM environments designed to elicit anxiety (Fig. 5), indicating an environment-specific anxious-like alteration following Dsp knockdown. The hippocampus plays functionally distinct roles in the dorsal and ventral regions (Fanselow and Dong, 2010). The dorsal region is involved in stress-induced learning and memory changes, and the ventral region contributes to stress responses and emotions such as anxiety. Because AAV was injected into the ventral DG area in this study (as described in the Methods section), Dsp in the ventral DG may play an anxiolytic-like role.

In animal models, chronic antidepressant treatment induces anxiolytic-like effects (Santarelli et al., 2003) and promotes the differentiation of newborn neurons in the DG (Wang et al., 2008; Ueno et al., 2019), while suppressing the expression of mature markers, including *Dsp,* in the DG (Fig. 1; Kobayashi et al., 2010; Imoto et al., 2017). The loss of Dsp function in the DG may lead to anxiolytic effects; however, surprisingly, anxiety-like behavior was observed (Fig. 5). Previous findings indicated that adult neurogenesis in the DG affects anxiety-like behaviors. For instance, transgenic mice in which adult neurogenesis was suppressed by the expression of the pro-apoptotic protein BAX in neural progenitor cells show increased anxiety-related behaviors (Revest et al., 2009). Therefore, suppression of adult neurogenesis in the DG by Dsp knockdown may promote anxiety-like behaviors.

Furthermore, acute activation of the ventral DG by channelrhodopsin in proopiomelanocortin (POMC)-Cre mice suppresses anxiety-like behavior in the EPM (Kheirbek et al., 2013). Because our findings showed that Dsp knockdown in the DG reduced the expression of the neuronal activity marker FosB (Fig. 2D), the suppression of neuronal activity in the DG by Dsp knockdown may also contribute to anxiety-like behavior.

Taken together, our findings demonstrate that Dsp in the DG regulates neuronal function and contributes to anxiolytic-like effects. This research uncovers an unexplored function for Dsp in the central nervous system and implies that Dsp in the DG may function as a regulator maintaining proper neuronal activation. Excessive antidepressant treatment can exacerbate neuronal activation, resulting in increased motor activity (Kobayashi et al., 2010; 2011). Considering the results of these studies and our findings by Dsp knockdown, reduction of Dsp expression by chronic antidepressants in the DG may function as a negative feedback mechanism for excessive neuronal activation rather than merely mediating the effects of antidepressants or anxiolytics.

### Limitations of the study

First, although we have proposed a role for Dsp in the nervous system, it is not clear what kind of complexes Dsp forms in neurons. In the future, it will be necessary to investigate what kind of factors Dsp binds to and forms complexes with. Second, we do not know how Dsp regulates neuronal activity and neurogenesis; elucidating the regulation of intracellular signaling by Dsp will improve our understanding of neuronal development and function. Third, it is not clear how Dsp in the DG contributes to emotional behaviors and whether Dsp in the DG contributes to learning and memory. The influence of Dsp in the DG on neural circuits involved in emotion needs to be elucidated. In addition, the role of Dsp in the dorsal DG needs to be investigated for its involvement in learning and memory. Finally, this study shows results in male mice only. It is possible that emotion-related behaviors such as anxiety have different phenotypes depending on sex. Future research should also investigate the role of Dsp in the female nervous system.

## Data availability statement

All datasets analyzed in the current study are available from the corresponding author upon reasonable request.

## Ethics statement

Animal use and procedures were conducted in accordance with the Tokyo University of Science regulations on animal care and use in research and were approved by the Animal Care and Use Committee of Tokyo University of Science (approval number K20009, K21007, and K22007).

## Author contributions

KO, NS, YN, YS, TT, KK, and KS designed the study, conducted the experiments, analyzed the data, and drafted the manuscript. ESN designed the study, analyzed the data, and drafted the manuscript. All the authors have read and approved the final manuscript.

## Acknowledgements

We would like to thank *Editage* (www.editage.jp) for the English language editing.

